# Bacterial isolates from CSF sample and their antimicrobial resistance patters in among under-five children suspected for meningitis in Dilla University Referral Hospital

**DOI:** 10.1101/2020.06.01.127456

**Authors:** Ephrem Awulachew, Kuma Diriba, Netsanet Awoke

**Author notes:** Corresponding author (Email =, Tell =).

## Abstract

**Introduction:** Bacterial meningitis is medical emergency that requires immediate medical attention. It is a cause of an estimated 288, 649 deaths worldwide per year, of which 94 883 death occur among Under-five children. Up to 24% of the survivors suffer from long-term sequelae such as epilepsy, mental retardation, or sensorineural deafness especially when the disease is contracted during early childhood.

**Objective:** the aim of this study was to assess bacterial isolates of CSF sample and their antimicrobial resistance patterns among under-five children in Dilla University Referral Hospital.

**Material and Methods:** Hospital based cross-sectional study design was used to collect clinical data and CSF sample from under-five children who was suspected for meningitis. Sediment of CSF sample was inoculated to Blood Agar plate, Chocolate Agar plate and Mackonkey Agar for bacterial isolation and identification. Chemical analysis and cytological analysis was also conducted based on standard operating procedures.

**Results:** From a total of 287 CSF sample cultured, causative bacteria were detected in 38 (13.2%). From culture positive cases the most frequent isolate was *Streptococcus pneumoniae* 13 (34.2%) followed by *Staphylococcus aureas* 7 (18.4%), *Neisseria meningitidis* 6 (16%) and *Escherichia coli* 6 (16%). *Haemophilus influenzae* b was isolated in 4 (10.5%) of children with meningitis. The other cause of meningitis was *Streptococcus agalactiae* which accounted (10.5%). Cryptococcus neoformans have been detected in 4 (1.9%) cases of meningitis. Of all bacterial isolates about 42.1% (16/38) bacterial isolates were multidrug resistant. About 38.5% of *S. pneumoniae* was multidrug resistance while about 33.3% *N. meningitis, 50% of H. influenzae, 57.1% of S. aureas* and 40% of *E. coli* showed multidrug resistance

**Conclusions:** High prevalence of bacterial meningitis and high rate of drug resistance were observed. *S. pneumoniae* was the leading cause of bacterial meningitis among under-five children.

## Introduction

Bacterial meningitis (BM) is an inflammation of the meninges, the protective linings of the brain and spinal cord caused by bacteria (1). In the past decades the epidemiology and treatment strategies for bacterial meningitis have significantly change (2). The introduction of conjugate vaccines in has resulted in reduction of *Haemophilus influenzae* type b, while conjugate pneumococcal and meningococcal vaccines have significantly reduced the burden of bacterial meningitis (3-5). Common etiologies of bacterial meningitis are; *Haemophilus influenzae* b, *Streptococcus pneumoniae, Neisseria meningitidis*, group B Streptococcus (GBS), and Listeria monocytogenes depending on age, sex, race, season and immunologic status of the child (6).

Bacterial meningitis is medical emergency that requires immediate medical attention (7, 8). Even with appropriate treatment, morbidity and mortality can be substantial (9, 10). Despite modern antibiotics and improved critical care, bacterial meningitis (BM) case fatality rates remain high (11, 12). It is a cause of an estimated 288, 649 deaths worldwide per year, of which 94 883 death occur among Under-five children (13, 14). Although effective vaccine reduces the incidence and effective modern antibiotics kill bacteria efficiently, mortality rates are still up to 34% in developing countries (15). BM usually causes serious complications such as brain damage, hearing loss, and ultimately death if not diagnosed and treated promptly (16, 17). Up to 24% of the survivors suffer from long-term sequelae such as epilepsy, mental retardation, or sensorineural deafness especially when the disease is contracted during early childhood (18).

A region of sub-Saharan Africa extending from Ethiopia in the East to Gambia in the West and containing fifteen countries and over 260 million people is known as the “meningitis belt” because of its high endemic rate of disease with superimposed, periodic, large epidemics. During epidemics, children and young adults are most commonly affected, with attack rates as high as 1,000/100,000 population, or 100 times the rate of sporadic disease (19, 20). The highest rates of endemic or sporadic disease occur in children less than 2 years of age. Case-fatality rates (CFRs) of suspected meningitis cases in the African meningitis belt were 4–26 % according to the country WHO weekly epidemiological record (21).

In Ethiopia, the disease has remained a serious health threat for the community especially among under five children. Bacterial meningitis occurs in Ethiopia particularly in the dry season from December to June and there were frequent meningococcal epidemics in different parts of the country (22, 23).

Another challenging epidemiologic trend is the emergence of antimicrobial resistance among meningeal pathogens. Drug resistance has been a reported all around worldwide (24, 25). However, the situation in developing countries like Ethiopia is particularly serious (7-8). Because of the absence of well-equipped bacteriological laboratories in Ethiopia no organized surveillance exists on drug resistance patterns among common bacterial isolates (26).

The early signs of meningitis are entirely nonspecific in the newborns and pediatrics patients unless the disease is positively excluded. Therefore, early diagnosis and appropriate antibiotic treatments are necessary to avoid further complications. Knowledge of the locally predominant meningeal pathogens and their sensitivity pattern is essential to plan for management bacterial meningitis and to minimize adverse outcomes. So the main aim of this study was to assess bacterial isolates of CSF sample and their anti-microbial resistance patterns among under-five children in Dilla University Referral Hospital, South Ethiopia.

## Materials and Methods

### Study area and Study period

The study was conducted in Dilla University Referral Hospital from February 1/2019 to march 30/2020. Dilla University referral is located in Dilla Town south of Ethiopia. It is located 356km far from Addis Ababa. The Hospital is the Referral Hospital in Gedeo zone, south of Ethiopia.

### Study design

Hospital based cross-sectional study design was used to collect clinical data and CSF sample for bacterial isolation.

### Study population

Under-five children who was suspected for meningitis and to who lumbar puncture was conducted during the study period was the study population.

### Eligibility criteria

All pediatrics patients (<5 years of age) that were clinically suspected for meningitis during the study period were included in the study. Meningitis was clinically suspected if the patient was presented with any of the following signs and symptoms: lethargic, vomiting, poor feeding, irritable, fever (≥ 38°C), and headache (above 2 year), meningeal irritation signs (Kerning or Brudzinski signs or neck stiffness, ≥ 1 year) on examination were considered critical for suspicion of meningitis (27).

### Sample size determination and Sampling technique

The sample size for this study was calculated using the formula for estimation of single population proportion considering prevalence of 30.6% from study conducted in Jimma, Ethiopia (28). So the total sample size was 287. Then the study participant was recruited consecutively until the required sample size reached using convenient sampling techniques.

### Data collection instrument and techniques

Interviewer administered questionnaire was developed through critical thinking and review of literatures and then translated to Amharic. Demographic information and clinical data o were collected using structured, interviewer administered questionnaire among the Children’s parents/guardian.

### Cerebrospinal Fluid sample collection and Identification of bacteria

#### CSF specimen collection

CSF sample is collected with lumbar puncture by physician and then immediately transported to Dilla University Microbiology Laboratory within 30 minutes for analysis. The collected CSF sample was drowned in to three consecutive test tubes according to standard of CSF analysis (29). CSF sample collected in the first test tube was used for chemical analysis (protein, glucose); the second test tube was used for microbiological analysis, while the third test tube was used for cytological analysis.

#### Processing CSF sample

Microbiological analysis of CSF sample: Once the CSF arrived in the Microbiology Laboratory, the volume of CSF available for analysis was noted. The sediment of centrifuged CSF was plated onto a blood agar plate (BAP) and a chocolate agar plate (CAP) and onto Mackonkey agar (MA) and was incubated aerobically and anaerobically at 37°c for 24-48 hrs. Purity plates (negative growth controls) were incubated after each new batch of produced media plates. Growth conditions were controlled by parallel incubation of the following control strains *S. pneumoniae ATCC 49619 and Escherichia coli ATCC 25922* obtained from Ethiopian Public Health Institutes in every batch of media preparation. Gram stain was conducted simultaneously using freshly prepared gram stain reagents. CSF sample was also examined by fluorescent microscope for diagnosis of tubercular meningitis and Indian ink preparation was used to diagnose for *Cryptococcal* meningitis.

Cytological examination of the CSF was conducted on the 3^rd^ test tube. CSF sample was noted for Turbidity, leukocytosis (usually of polymorphonuclear (PMN) leukocytes); WBC counts >10 cells/mm. Chemical analysis of CSF sample was conducted using spectrophotometer to measure glucose and protein concentration of CSF sample at 546nm wavelength.

#### Isolation and identification of bacteria

After overnight incubation of the inoculum at 37 °C, BAP, CAP and MA culture plates were examined for colonies. All positive CSF cultures was identified and characterized on the basis of morphology, cultural characteristics, and biochemical testing and serogroup-specific antisera.

Identification of *Neisseria meningitidis* (*N. meningitides*) by their round, moist, glistening, and convex colonies on BAP or CAP, Gram negative diplococcic, oxidase positive, acid production in glucose and maltose and not in sucrose or lactose. Then the serogroup of *N. meningitidis* were identified by using serogroup-specific antisera (Bio-Rad., Pastorex^™^ Meningitis, lot No:-64247667). *Haemophilus influenzae* (*H. influenzae*) isolate was identified by taking colorless-to-grey, opaque colonies grown in the presence of X and V factor on chocolate agar was tested for oxidase positivity and finally it was serotyped using *H. influenza* type b serogroup-specific antisera (Bio-Rad., Pastorex^™^ Meningitis). *Streptococcus pneumonia* (*S. pneumonia*) was identified by their grey moist flattened and depressed center colonies with a zone of alpha-hemolysis on BAP, Gram reaction, catalase negative, optochin sensitivity (>14mm), and bile solubility. Then the *S. pneumonia* isolate was serotyped using serogroup-specific antisera (Bio-Rad., Pastorex^™^ Meningitis, lot No:-64247667). *Streptococcus agalactiae* (*S. agalactiae*) was identified by characteristic colonies of translucent grey colonies, with a complete ß hemolysis on BAP, Gram reaction, catalase negative, and esculin negative. Then the isolates were serotyped using serogroup-specific antisera (Bio-Rad., PastorexTM Meningitis, lot No:- 64247667).

Other gram positive and gram negative organisms were identified by series of biochemical tests like catalase test, coagulase test, Triple Suger Iron slant agar (TSI) (Oxoid), indole test and urase test (Oxoid) and Sulfide-Indole-Motility (SIM).

#### Antibiotic sensitivity test

The antibiotic resistance patterns of bacterial isolates against commonly used antibiotics was conducted on Mueller Hinton agar (MHA) (Oxoid) and incubated at 37°c; for 24 hrs. The standard disk diffusion technique of modified Kirby-Bauer method was used as recommended by European Committee on Antimicrobial Susceptibility Testing (EUCAST) (30). For the susceptibility testing the following eight antimicrobial drugs and concentrations was used: penicillin G (10μg), chloramphenicol (30 μg), cefotaxime (5μg), ciprofloxacin (5μg), Ceftazidime (10μg), ceftriaxone (30μg), Vancomycin (15μg) and rifampicin. Multidrug resistance was considered when résistance to 2 or more drugs belonging to different classes of antibiotics tested (31, 32).

### Study variables

#### Dependent/outcome variables

Prevalence of *bacterial meningitis*

Patterns of drug resistance

#### Independent/explanatory variables

Socio-demographic variables like age and sex

Clinical data (fever, irritability, neck stiff, poor feeding, light intolerance)

### Data quality assurance

Training was given to laboratory assistant and data collectors for two days on the procedures of data collection and handling of collected data. The collected data was checked for completeness at the end of data collection. During laboratory analysis of CSF cultures, standard operating procedures were followed. To maintain the quality of the result, CSF sample grossly contaminated with blood, sample not received within two hours of collection, and sample not in appropriate transport medium were rejected. Culture media was prepared and sterilized based on SOPs. Then the sterility of culture media was checked by parallel incubation of un inoculated plate in every newly prepared culture plates and observed for bacterial growth. Control strains *S. pneumoniae* ATCC 49619 and *E. coli* ATCC 25922 that were obtained from Ethiopian Public Health Institute was used as a quality control during CSF cultures, biochemical tests and antimicrobial susceptibility testing.

### Data analysis

The Collected data was coded, entered and cleaned using Epi-Data version 3.02 and transported to SPSS version 20 for analysis. Descriptive statistics such as frequency, percentage and cross tabulation was used to present the findings. Chi-square was performed to evaluate whether the variables have significant association with the outcomes of interest at 95% confidence interval.

### Ethical consideration

Institutional ethical clearance was obtained from Dilla University Health Research Ethics Review Committee. During data collection each child’s parent/guardian was informed about the aim of the study. Written consent was obtained before the start of data collection. Samples with positive culture results were communicated to physicians in order for patients to get treatment according to drug susceptibility results of isolates.

## Results

### Socio-demographic characteristics

About 287 suspected pediatric meningitis cases were examined for bacterial isolate using culture techniques in Dilla University Microbiology Laboratory. Of whom 163 (56.8%) were male and 124 (43.2%) were female. The mean age of the participant children were 2.6+ 0.7 year. Majority 169 (58.9%) of the patients were from Rural area. Majority of the children were in the age group of 13-59 months of age (41.5%) (Table 1).

**Table: 1.** Socio demographic characteristics and culture positivity

| Socio demographic characteristics of the patients |  | Number (n) | Culture positive | Culture negative | Cryptococcus neoformans |
| --- | --- | --- | --- | --- | --- |
| Sex | Male | 163 | 21 | 139 | 3 |
|  | Female | 124 | 17 | 106 | 1 |
| Age | 0-28days | 64 | 10 | 54 | 0 |
|  | 1-12 months | 104 | 14 | 89 | 1 |
|  | 13-59 months | 119 | 14 | 102 | 3 |
| Residence | Urban | 169 | 25 | 140 | 4 |
|  | Rural | 118 | 13 | 105 | 0 |

In about 95 (33.1%) and 83 (28.9%) patients, CSF samples were collected within 12 hours and 24 hours after empirical treatments respectively. About 109 (38%) of meningitis suspect pediatrics patients started empirical treatment immediately after CSF samples collected. The characteristics of 73 (25.4%) CSF samples collected were purulent with a visible turbidity and 11 (3.8%) CSF samples were xanthochromic, while the remaining 203 (70.7%) CSF samples were clear in appearance. Pleocytosis (>50 cells/μl) were observed in 164 (57.1%) of CSF sample with a median white blood cell count of 340 cells/μl. From 164 patients with pleocytosis, granulocytosis (predominantly neutrophil) seen in 97 (59.1%) of patients while 67 (40.9%) patients had lymphocytosis. Spectrophotometric measurements of CSF protein and glucose showed that 183 (63.8%) CSF sample had normal protein with a mean protein level of 47.8 mg/dl ± 9.6 mg/dl and 87 (30.3%) CSF sample had elevated protein with a mean protein concentration of 79 mg/dl ±11.6 mg/dl. About 201 (70%) CSF sample had normal glucose concentration with a mean glucose centration of 64 mg/dl ± 6.9 mg/dl and 65 (22.6%) had reduced glucose concentration with a mean concentration of 32 mg/dl ± 4.7 mg/dl.

### Clinical data

Clinical sign and symptoms of children with suspected meningitis was collected using standard questionnaire by trained nurses. According to the results of clinical data obtained fever was demonstrated in 87% of cases, while irritability, rash and poor feeding seen in 67%, 26% and 91% respectively (Figure: 1).

**Figure: 1.**
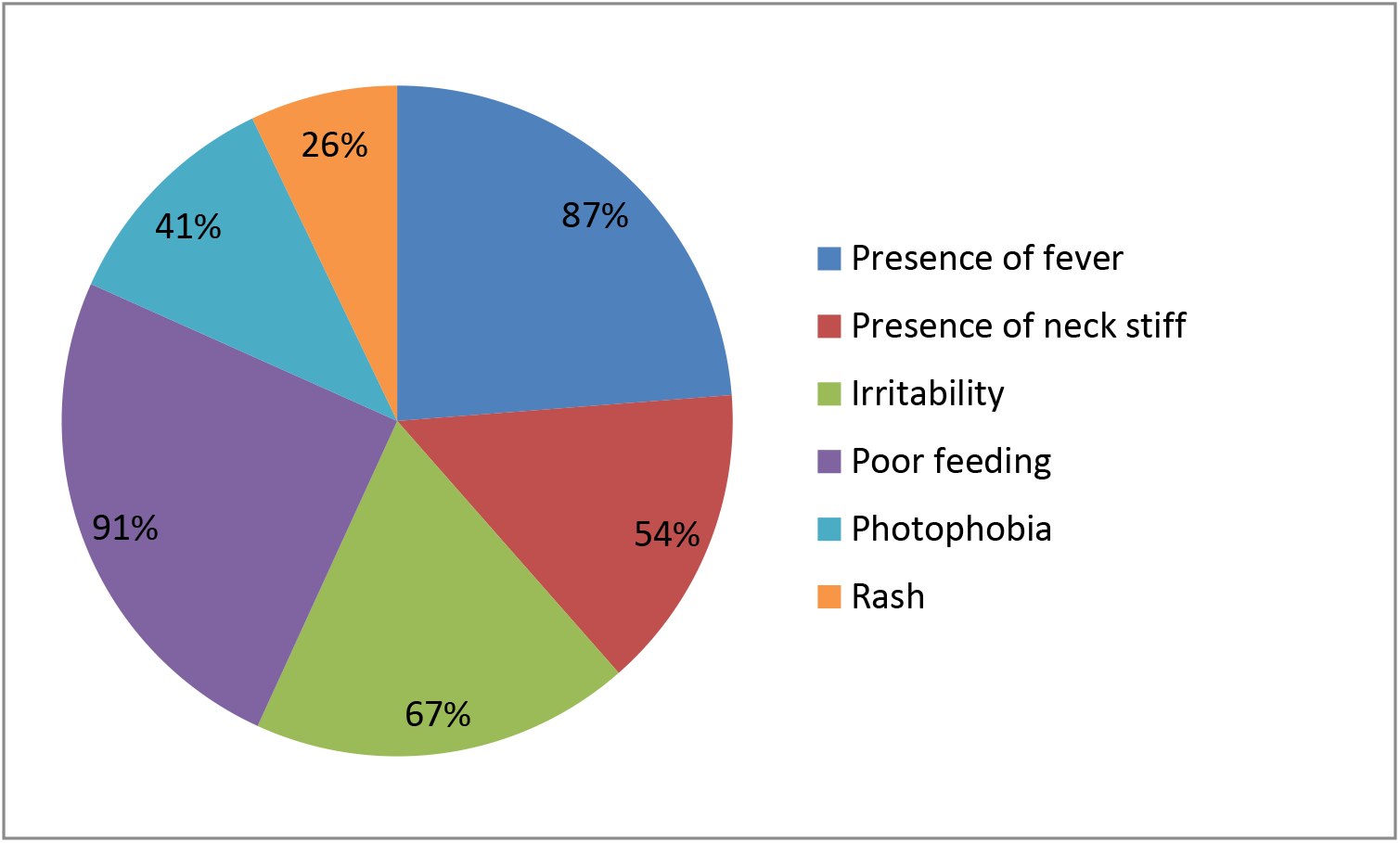
Clinical features of children with bacterial meningitis

### Etiology of meningitis

From a total of 287 CSF sample cultured, causative bacteria were detected in 38 (13.2%). This makes the overall prevalence of bacterial meningitis 13.2% (38/287). In present study, gram stains were performed directly from CSF sample and the yield was 7.3% (21/287). From culture positive cases the most frequent isolate was *Streptococcus pneumoniae* 13 (34.2%) followed by *Staphylococcus aureas* 7 (18.4%), *Neisseria meningitidis* 6 (16%) and *Escherichia coli* 6 (16%). *Haemophilus influenzae b* was isolated in 4 (10.5%) of children with meningitis. The other cause of meningitis was *S. agalactiae* which accounted 4 (10.5%). *Cryptococcus neoformans* have been detected in 4 (1.9%) cases of meningitis (Table 2).

**Table: 2.**
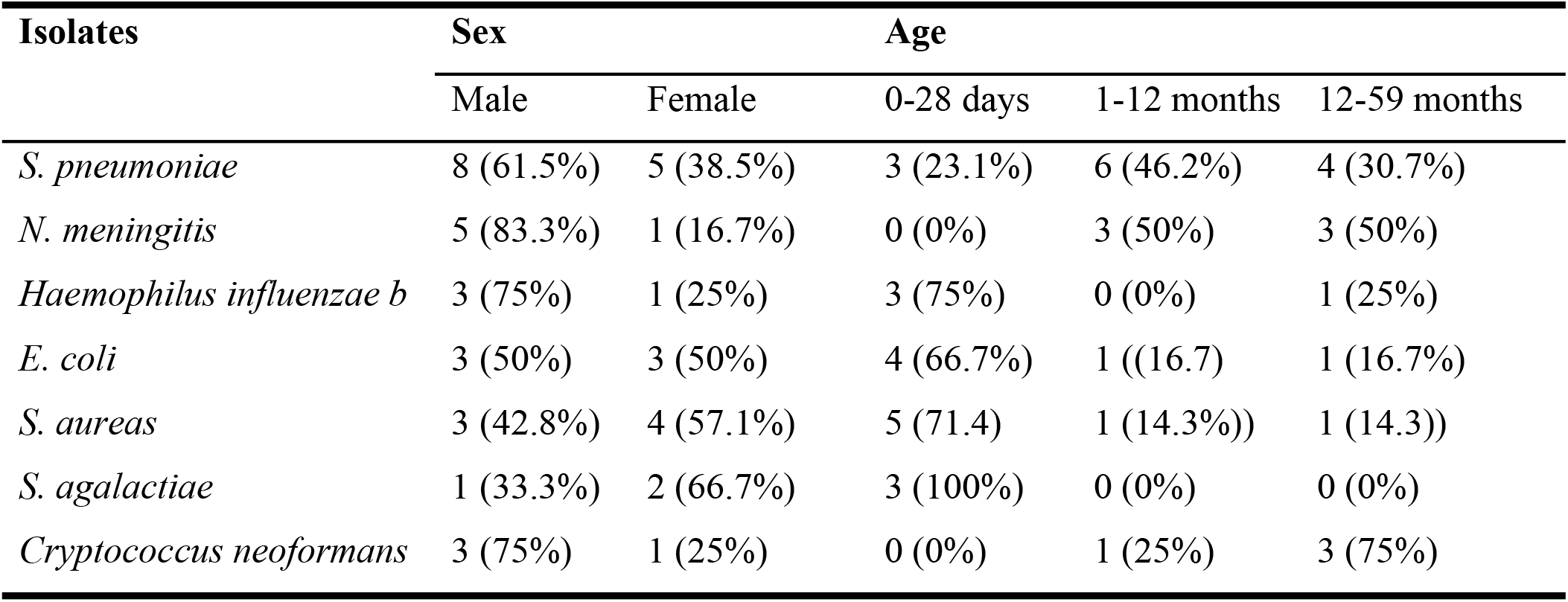
Microbial isolates of CSF samples from under-five children suspected for meningitis

### Serotypes of the isolate

Based on serological characterization the most frequent *Neisseria meningitidis* were Y/W135, B and A with a prevalence of 50% (2/4), 25% (1/4), and 25% (1/4) respectively. All four if *H. influenzae* isolates was type b.

There is no statistical significant association with isolated bacteria and gender of the patients (P = 0.521). Majority of the isolate (43.5%) of bacterial meningitis occurred in 1-12 months and 13-59 months age groups. Regarding statistical association of bacterial isolate, *S. agalactiae* (group B streptococci (GBS)) were frequently found in children with the age of 0-28 days (*P* = 0.010). *H. influenzae b* infections showed statistical association with the age of the children, where it significantly affect children with the age of 0-28 days (*p* < 0.001). On the other hand *N. meningitidis* was significant causes of meningitis among children with the age range of 1-5 years (OR= 1.8, *p* = 0.021).

### Anti-microbial resistance

Antimicrobial susceptibility tests indicated that the resistance rates of *S. pneumoniae* isolates to penicillin G, chloramphenicol, cefotaxime, ciprofloxacin, and ceftriaxone were 38.5%, 38.5%, 30.8%), 15.3%, and 15.4% respectively. The drug susceptibility test result of *N. meningitidis* to penicillin G was 33.3%, while resistance to chloramphenicol, ceftazidime, ciprofloxacin, and ceftriaxone were 16.7% each. *N. meningitidis* also showed 33.3% resistance to rifampicin. The resistance rate of *H. influenzae* to penicillin G, chloramphenicol, cefotaxime, ciprofloxacin, and ceftriaxone were 50%, 25%, 50%, 25% and 20% respectively. The rate of resistance of *E. coli* to chloramphenicol, ciprofloxacin and ceftriaxone were 60%, 40% and 20% respectively. Antimicrobial resistance of *S. aureas* to penicillin G, chloramphenicol, vancomycin and ceftriaxone were 28.6%, 57.1%, 14.3% and 14.3% respectively. *S. agalactiae* showed a resistance of 33.3% to penicillin G and rifampicin (Table 3).

**Table: 3.** Antimicrobial resistance patterns of the isolates CSF sample

| Drugs |  | *S.*<br>*pneumoniae* | *N.*<br>*meningitis* | *H.*<br>*influenzae* | *E. coli* | *S. aureas* | *S. agalactiae* |
| --- | --- | --- | --- | --- | --- | --- | --- |
| **Penicillin G** | **S** | 8 (61.5%) | 4 (66.7%) | 2 (50%) | 2 (40%) | 3 (42.8%) | 2 (66.7%) |
|  | **I** | 0 (0%) | 0 (0%) | 0 (0%) | 0 (0%) | 2 (28.6%) | 0 (0%) |
|  | **R** | 5 (38.5%) | 2 (33.3%) | 2 (50%) | 3 (60%) | 2 (28.6%) | 1 (33.3%) |
| **Chloramphenicol** | **S** | 7 (53.8%) | 4 (66.7%) | 3 (75%) | 2 (40%) | 3 (42.9%) | 2 (66.7%) |
|  | **I** | 1 (7.7%) | 1 (16.7%) | 0 (0%) | 0 (0%) | 0 (0%) | 0 (0%) |
|  | **R** | 5 (38.5%) | 1 (16.7%) | 1 (25%) | 3 (60%) | 4 (57.1%) | 1 (33.3%) |
| **Cefotaxime** | **S** | 9 (69.2%) | 4 (66.7%) | 2 (50%) | 5 (100%) | 4 (57.1%) | 3 (100%) |
|  | **I** | 0 (0%) | 2 (33.3%) | 0 (0%) | 0 (0%) | 0 (0%) | 0 (0%) |
|  | **R** | 4 (30.8%) | 0 (0%) | 2 (50%) | 0 (0%) | 3 (42.9%) | 0 (0%) |
| **Ciprofloxacin** | **S** | 10 (77%) | 5 (83.3%) | 3 (75%) | 3 (60%) | 5 (71.4%) | 2 (66.7%) |
|  | **I** | 1 (7.7%) | 0 (0%) | 0 (0%) | 0 (0%) | 0 (0%) | 0 (0%) |
|  | **R** | 2 (15.3%) | 1 (16.7%) | 1 (25%) | 2 (40%) | 2 (28.6%) | 1 (33.3%) |
| **Ceftazidime** | **S** | 9 (69.2%) | 5 (83.3%) | 3 (75%) | 4 (80%) | 5 (71.4%) | 2 (66.7%) |
|  | **I** | 2 (15.4%) | 0 (0%) | 0 (0%) | 0 (0%) | 0 (0%) | 0 (0%) |
|  | **R** | 2 (15.4%) | 1 (16.7%) | 1 (25%) | 1 (20%) | 2 (28.6%) | 1 (33.3%) |
| **Ceftriaxone** | **S** | 11 (84.6%) | 5 (83.3%) | 3 (75%) | 4 (80%) | 6 (85.7%) | 3 (100%) |
|  | **I** | 0 (0%) | 0 (0%) | 0 (0%) | 0 (0%) | 0 (0%) | 0 (0%) |
|  | **R** | 2 (15.4%) | 1 (16.7%) | 1 (25%) | 1 (20%) | 1 (14.3%) | 0 (0%) |
| **Vancomycin** | **S** | 10 (77%) | 5 (83.3%) | 3 (75%) | 4 (80%) | 6 (85.7%) | 3 (100%) |
|  | **I** | 2 (15.3%) | 0 (0%) | 0 (0%) | 0 (0%) | 0 (0%) | 0 (0%) |
|  | **R** | 1 (7.7%) | 1 (16.7%) | 1 (25%) | 1 (20%) | 1 (14.3%) | 0 (0%) |
| **Rifampicin** | **S** | 9 (69.2%) | 4 (66.7%) | 2 (50%) | 2 (40%) | 2 (28.6%) | 2 (66.7%) |
|  | **I** | 1 (7.7%) | 0 (0%) | 0 (0%) | 0 (0%) | 0 (0%) | 0 (0%) |
|  | **R** | 3 (23.1%) | 2 (33.3%) | 2 (50%) | 3 (60%) | 5 (71.4%) | 1 (33.3%) |

**Table 4.** Antimicrobial resistance pattern of bacterial isolates from CSF samples

| Organisms | MDR level of bacterial isolates | | | | | Total MDR isolates (≥ R2) |
| --- | --- | --- | --- | --- | --- | --- |
| | R0 | R1 | R2 | R3 | >R4 | ≥ R2 |
| *S. pneumoniae* | 3 | 5 | 1 | 2 | 2 | 5 |
| *N. meningitidis* | 1 | 3 | 0 | 1 | 1 | 2 |
| *H. influenzae b* | 0 | 2 | 0 | 0 | 2 | 2 |
| *E. coli* | 1 | 2 | 0 | 1 | 1 | 2 |
| *S. aureas* | 1 | 2 | 1 | 1 | 2 | 4 |
| *S. agalactiae* | 2 | 1 | 1 | 0 | 0 | 1 |

## Discussion

In this study the overall prevalence of bacterial meningitis among under-five children was 13.2% which is lower than study conducted in Jimma, Ethiopia that reported prevalence of 30.6% from 85 CSF sample analyzed (28). This difference could be due to small size of study conducted in Jimma. The other possible causes of the difference could be time of CSF sample collection, in this study about 62% of CSF sample were collected after empirical antimicrobial treatment while study conducted in Jimma reported that CSF samples were collected after treatment in about 52.9% patients (28). CSF sample collection after prior antimicrobial treatment could obviously contribute to culture negative results. The overall prevalence of bacterial meningitis of the current study was also lower than study conducted in Bahr Dar that reported a prevalence of 69.8% (33). This difference could be due to difference diagnostic method. For diagnosis of bacterial meningitis, study conducted in Bahr Dar mainly dependent of clinical diagnosis and Gram stain. On the other hand the overall prevalence of bacterial meningitis was comparable with study conducted in Addis Ababa, Ethiopia, that reported a prevalence of 14.1% (34).

In the present study the predominant cause of bacterial meningitis among under-five children was *S. pneumoniae* (34.2%), this is similar with study conducted in China where *S. pneumoniae* was reported to be the predominant isolate (35). This finding of the present study could be supported by study conducted in Gonder, in Addis Ababa and in Jimma which reported that the predominant isolates was *S. pneumoniae* 70%, 35.3% and 50% respectively (28, 36, 37).

In the current study from culture proven cases the prevalence of *N. meningitidis* was 16%. This higher than study conducted in china (3.8%) (38) and Addis Ababa (3.2%) (34). *Haemophilus influenzae* b was isolated in 4 (10.5%) of children with meningitis. This finding is comparable with study conducted in china that reported a prevalence rate of *H. influenzae* b (9.5%) (35) and it is also consistent with study conducted in Jimma, Ethiopia which demonstrated 11.5% prevalence of *H. influenzae* b from culture proven cases (28). On the other hand, this finding is slightly higher than prevalence of *H. influenzae* b reported in study conducted in India (1.4%). This difference could be due to large sample size of study conducted in India and as well as socio-economic difference of the countries (39).

*Escherichia coli* (66.7%) and *S. agalactiae* (100%) were the predominant organisms causing bacterial meningitis in children with the age of 0-28 days. This is consistent with study conducted in China (38). These could be due to the fact that it will be accidentally inoculated to newborns from birth canal during delivery.

Regarding antimicrobial susceptibility test, majority of the isolates were resistant to commonly used antibiotics such as penicillin G, chloramphenicol, ciprofloxacin and ceftriaxone. In our study antimicrobial susceptibility tests indicated that about 38.5% isolates of *S. pneumoniae* were resistant to penicillin G which is in line with other study in Gonder, Ethiopia (43%) (40) and in Nigeria (41). On the other hand penicillin resistance rate of *S. pneumoniae* in this study was slightly lower than study conducted in China that reported penicillin resistance rate of *S. pneumoniae* was 68.8% (35). This high rate of penicillin resistance in all reported studies indicates that the need for continued monitoring for penicillin resistance in pneumococcal isolates due to frequent use of the drug.

On the other hand more than half of *E. coli* (60%) was resistant to penicillin G in this study, which in line with study conducted in china which showed more than half of *E. coli* showed resistance to penicillin (42). In the present study *E. coli* showed the highest resistance to penicillin G (60%) followed by *H. influenzae* b (50%) and *S. pneumoniae* (38.5%). One isolates of *H. influenzae* b showed lower rate of resistance to ciprofloxacin and ceftriaxone. This is in line with study in china that reported lower rate resistance to ciprofloxacin and ceftriaxone (35). Unlike study conducted in Gonder where no organism was found to be resistant to ciprofloxacin, in the present study *E. coli, S. agalactiae* and *S. aureas* showed 40%, 33.3% and 28.6% resistance to ciprofloxacin while *H. influenzae, N. meningitidis* and *S. pneumoniae* showed a resistance of 25%, 16.7% and 15.3% respectively. This could be due to gradual rise in multi-drug resistant species bacteria.

The antimicrobial resistance rate of *N. meningitis* to chloramphenicol and penicillin were 16.7% and 33.3% respectively. This is similar with study conducted in Gonder, Ethiopia that reported the *N. meningitidis* was found to be resistant to chloramphenicol 30%, and penicillin 20% (40).

High rate of multi drug resistant isolates were also observed. Of all bacterial isolates, 42.1% of the bacterial isolates were multidrug resistant. This finding is comparable with study conducted in Gonder (40). About 38.5% of *S. pneumoniae* was multidrug resistance while about 33.3% *N. meningitides*, 50% of *H. influenzae*, 57.1% of S. aureas and 40% of *E. coli* showed multidrug resistance.

## Limitation of the study

Despite good outcome we acknowledge few limitations. The first limitation was that this study considered only CSF sample while additional blood culture could be important. The small sample size was the other limitation of this study.

## Conclusion

In the present study high prevalence of bacterial meningitis was detected. From culture proven meningitis, *S. pneumoniae* was the leading cause of bacterial meningitis among under-five children. *N. meningitidis* serotype Y/W135 was the most common strain isolated from CSF sample. *H. influenzae* type b was an important cause of bacterial meningitis and relatively common among children with the age of 0-28 days. High rate of resistance were observed for most of the isolates of CSF sample. About 38.5% of *S. pneumoniae* was multidrug resistance while about 33.3% *N. meningitis, 50% of H. influenzae, 57.1% of S. aureas* and 40% of *E. coli* showed multidrug resistance.

## Declarations

### Conflicts of interest

The Authors have no conflicts of interests to declare.

### Availability of data and materials

All the datasets generated and analyzed during the review are included in this article.

### Author’s contribution

E. A, K. D, and N.A designed the study, extracted, critically reviewed and analyzed data and wrote the first draft of the manuscript, and approved the manuscript.

### Funding source

This study was funded by Dilla University Research Dissemination Office (DURDO)

### Consent for publication

Not applicable.

## Acknowledgements

First of all, I would like to thank Dilla University for opening this opportunity for researcher to come with problem solving project ideas and scientific questions and we also acknowledge Dilla University Research Dissemination Office for finding this study.

My acknowledgement also goes to librarians for their support in providing me journals and relevant literatures kindly.

